# Hyperintense signals in cerebral blood flow maps acquired with pseudo-continuous arterial spin labeling MRI in mice

**DOI:** 10.64898/2025.12.02.691929

**Authors:** Xiuli Yang, Yuguo Li, Adnan Bibic, Zhiliang Wei

**Affiliations:** Russell H. Morgan Department of Radiology and Radiological Science, Johns Hopkins University School of Medicine, Baltimore, Maryland 21205, USA; F. M. Kirby Research Center for Functional Brain Imaging, Kennedy Krieger Research Institute, Baltimore, Maryland 21205, USA

**Keywords:** hyperintensity, vascular suppression, bolus arrival time, crusher gradient, laminar flow, intravoxel dephasing

## Abstract

**Background and Purpose:** Pseudo-continuous arterial spin labeling (pCASL) MRI is a widely used, noninvasive, contrast-agent-free technique for measuring cerebral blood flow (CBF) and assessing vascular dysfunction across diverse clinical settings and murine disease models. In practice, arterial-transit artifacts that generate hyperintense signal in CBF maps warrant careful consideration. While these effects are well characterized in humans, they are less well understood in mice owing to the marked interspecies physiological differences.

**Methods:** To address this knowledge gap, we systematically characterized pCASL hyperintense signal as a function of post-labeling delay (PLD) and crusher-gradient strength in mice. Numerical simulations were also performed to validate the experimental findings.

**Results:** We found that hyperintense signals in mice extend to arteries, major veins, and ventricular structures (e.g., choroid plexus). Such a pattern was different from human pCASL images, where hyperintense signals are predominantly present in arteries. Statistical analyses supported a PLD of 500 ms as a pragmatic balance between detection sensitivity and suppression of vascular contamination. Additional experiments and numerical simulations showed that, within the tested range, stronger crusher gradients provided little extra vascular suppression—primarily because large vessel calibers relative to small voxels limit intravoxel phase dispersion. These findings refine the interpretation of murine pCASL signals and facilitate more accurate perfusion imaging in preclinical pathophysiological studies.

## 1. Introduction

Pseudo-continuous arterial spin labeling (pCASL) MRI provides noninvasive, contrast-free, and reproducible measurements of cerebral blood flow (CBF) by using magnetically labeled arterial water as an endogenous tracer [1, 2]. Owing to its fully noninvasive design, pCASL is well suited for longitudinal studies in mice and humans, enabling within-subject tracking of disease progression and therapeutic response [1]. Voxel-wise CBF maps are sensitive to regional hypoperfusion and hyperemia associated with cerebrovascular pathology, including neuroinflammation and neurodegeneration [3, 4]. Emerging evidence indicates that pCASL-derived CBF is a quantitative biomarker that complements structural and diffusion MRI [5], supports mechanistic neurovascular investigations [6], and provides clinically translatable endpoints for therapeutic interventions [7].

Hyperintense voxels in pCASL-based CBF maps primarily reflect intravascular contamination, i.e., labeled arterial water that has not yet exchanged with tissue, leading to overestimation of CBF in voxels dominated by blood rather than capillary exchange [1, 8]. In human pCASL studies, a sufficiently long post-labeling delay (PLD) is typically chosen to mitigate this arterial-transit artifact [1], but in cases with very long bolus arrival (arterial transit) times the optimal PLD becomes difficult to select without sacrificing detection sensitivity [9]. Bipolar crusher gradients provide an alternative means of vascular suppression [10, 11], yet they lengthen the echo time (TE) and can impose a substantial sensitivity penalty at ultrahigh field strengths typical of preclinical scanners (e.g., 7.0–11.7 T). Moreover, interspecies differences in brain physiology between humans and mice can further complicate these effects, contributing to the persistence and variability of hyperintensity in pCASL-derived CBF maps.

There is an unmet need to characterize the hyperintense signals observed in murine pCASL MRI. Addressing this gap will improve interpretation of pCASL readouts in mice and facilitate their application to pathophysiological studies with animal models. Accordingly, we systematically examined the behavior of hyperintense voxels in pCASL-based CBF maps across PLDs and evaluated the efficacy of vascular-crushing gradients for suppressing intravascular signal. Numerical simulations were performed to corroborate and interpret the experimental findings. Experiments and simulations revealed that hyperintense voxels persist in arteries and major draining veins across practical PLDs in mice, and crusher gradients provide limited incremental suppression relative to PLD tuning.

## 2. Methods

### 2.1 General

Experimental protocols for this study were approved by the Johns Hopkins Medical Institution Animal Care and Use Committee and conducted in accordance with the National Institutes of Health guidelines for the care and use of laboratory animals. Data reporting complied with the ARRIVE 2.0 guidelines. All procedures were carefully designed to minimize discomfort and stress to the animals. Mice were housed in a quiet environment under a 12-h light/dark cycle with ad libitum access to food and water. A cohort of 24 C57BL/6 mice (age: 33-37 weeks; body weight: 24-43 grams; 12 females, 12 males) was used. Since MRI is a non-invasive technique, mice were used in multiple experimental sessions. For mice scanned on the same day, the experimental order was randomized according to a previously reported scheme [12].

### 2.2 MRI

An 11.7 T Bruker Biospec system (Bruker, Ettlingen, Germany) with a horizontal bore and actively shielded pulse field gradient (maximal intensity: 0.74 T/m) was used for imaging. Data were acquired using a 72-mm quadrature volume resonator as the transmitter and a four-element (2 × 2) phased-array coil as the receiver. B0 homogeneity across the mouse brain was optimized using global shimming (up to the second order) based on a subject-specific pre-acquired field map. Inhalational isoflurane was used as the anesthetic following a previously reported scheme [13, 14]. Respiration was observed during the experiments using an MRI-compatible monitoring system (SA Instruments, Stony Brook, USA). Isoflurane dose was adjusted as needed to maintain a respiration rate of 70 – 120 breaths per minute. Experiments were terminated if respiration rates dropped below 50 breaths per minute for more than two minutes. A temperature-controlled water circulation system embedded in the scanner was used to maintain body temperature in the mice.

### pCASL scans in varying PLD values (N = 20)

A two-scan pCASL scheme developed to improve the robustness of arterial spin labeling against magnetic-field inhomogeneity was utilized in this study [15]. A pre-scan was first performed to optimize the phase offsets of labeling pulses for both control and label scans [16]. Subsequently, pCASL scans focusing on perfusion imaging were performed under the following parameters [17]: repetition time (TR) = 5000 ms; echo time (TE) = 11.8 ms; field of view = 15 × 15 mm²; slice thickness = 0.75 mm; interslice gap = 0.25 mm; matrix size = 96 × 96; number of slices = 12; receiver bandwidth = 300 kHz; labeling duration = 1800 ms; inter-labeling pulse delay = 1.0 ms; labeling pulse width = 0.4 ms; labeling plane thickness = 1.0 mm; labeling pulse flip angle = 40°; mean gradient = 0.01 T/m (label scans) and 0 T/m (control scans); post-labeling delay (PLD) = 25, 50, 100, 150, 200, 250, 300, 350, 400, 500, 600, 700, 1000, 1300, 1600, and 2000 ms; signal average = 12; and scan duration = 4.0 min using a two-segment spin-echo echo-planar-imaging (EPI) acquisition.

Time-of-flight (TOF) angiography was used to visualize the major cerebral arteries and veins in mice. Experimental parameters were: TR/TE = 20/2.7 ms; field of view = 15 × 15 mm²; slice thickness = 0.75 mm; interslice gap = 0.25 mm; matrix size = 256 × 256; number of slices = 12 (axial orientation); receiver bandwidth = 125 kHz; and scan duration = 0.8 min.

A cohort of ten mice (five females) was used for the pCASL and TOF scans. In a separate cohort of ten mice (five females), a spin echo sequence was performed with a long TE to visualize the cerebrospinal fluid (CSF) under the following parameters: TR/TE = 2500/150.0 ms; field of view = 15 × 15 mm²; slice thickness = 0.75 mm; interslice gap = 0.25 mm; matrix size = 96 × 96; number of slices = 12 (axial orientation); receiver bandwidth = 75 kHz; and scan duration = 4.0 min.

The processing of pCASL data followed established procedures [5]. Pairwise subtraction between control (M_CtT_) and label images (M_lbl_) was first performed to yield a difference image, which was then divided by an M_0_ image (obtained by scaling the control image [15]) to provide a perfusion index image: CBF_index_ = (M_CtT_ - M_lbl_)/M_O_. The CBF index maps were co-registered and normalized to a mouse brain template [18] and resized to recover the original acquisition resolutions. A histogram analysis on the mean CBF index maps was performed, followed by Gaussian fitting to derive a threshold for identifying hyperintense voxels. TOF and CSF images underwent similar processing to align anatomical landmarks across mice. Regions of interest (ROIs) were drawn on the CBF index image to encompass the cortex, dorsal cerebellar artery (DCA), posterior cerebral artery (PCA), posterior communicating artery (PComA), azygos pericallosal artery (azPA), anterior cerebral artery (ACA), transverse sinus (TS), confluence of sinuses (CS), great vein of Galen (VG), thalamostriate vein (TSV), superior sagittal sinus (SSS), and lateral ventricle (LV) by reference to the Allen Brain Atlas (https://atlas.brain-map.org/) and a mouse vasculature atlas [19]. ROIs were filtered with a hyperintensity mask (derived from the threshold) to isolate hyperintense voxels and were grouped into four categories: (a) tissue (cortex); (b) arteries (DCA, PCA, PComA, azPA, ACA); (c) veins (TS, CS, VG, TSV, SSS); and (d) ventricle (LV). Voxel-wise CBF indices within each ROI were averaged to represent the corresponding regions.

### pCASL scans with varying vascular suppression gradients (N = 4)

Four mice (2 females) were tested in Study 3. The pCASL scan in Study 1 was used. Crusher gradients straddling the refocusing pulse of the spin-echo EPI module were applied to suppress vascular signals. Crusher gradient and refocusing pulse durations were 1.5 ms and 1.7 ms, respectively. A series of triple-axis crusher gradient strengths, i.e., 0, 3.7, 7.4, 14.8, 22.2, 29.6, and 37.0 G/cm, were applied to evaluate vascular suppression performance. These gradient combinations corresponded to velocity encoding (VENC) values of infinite, 12.4, 6.2, 3.1, 2.1, 1.6, and 1.2 cm/s, respectively. The pCASL dataset was processed as described in Study 2.

Three simulations were performed to investigate the effects of artery diameter, voxel size, and flow pattern on the performance of vascular suppression as follows. (Simulation 1) Effect of artery diameter and voxel size: peak flow velocity = 10 cm/s (laminar flow assumed), crusher gradient strength = 20 Gauss/cm, artery diameter = 100, 120, 140, …, 400 µm, voxel size = 1, 5, 10, 15, …, 300 µm. (Simulation 2) Effect of flow pattern: peak flow velocity = 10 cm/s, crusher gradient strength = 20 Gauss/cm, voxel size = 100 µm, artery diameter = 100, 120, 140, …, 400 µm, laminar fraction = 0, 0.05, 0.10, …, 1 (fraction of laminar flow competed by the plug flow; for example, 0.50 indicated 50% laminar flow with 50% plug flow). (Simulation 3) Effect of crusher gradient strength: peak flow velocity = 10 cm/s (laminar flow assumed), artery diameter = 300 µm, crusher gradient strength = 0, 5, 10, …, 70 Gauss/cm, and voxel size = 1, 5, 10, 15, …, 300 µm. The crusher gradient and refocusing pulse durations of 1.5 ms and 1.7 ms were used in all simulation.

### 2.3 Data processing

All data processing was performed using custom MATLAB scripts (MathWorks, Natick, MA).

### 2.4 Statistical analyses

Linear mixed-effects models were used to examine the dependence of regional CBF index on PLD. We fit two specifications: CBF_index_∼PLD + (1|Mouse) and CBF_index_ ∼ PLD * Region + (1|Mouse), where PLD was treated as a continuous predictor, Region as a categorical factor, and a random intercept was included for Mouse. Linear mixed-effects models were also used to examine the dependence of regional CBF index on vascular-crushing gradient strength (Grad). We fit two formula: CBF_index_∼Grad + (1|Mouse) and CBF_index_ ∼ Grad * Region + (1|Mouse), treating Grad as a continuous predictor. Linear mixed-effects models were fit by restricted maximum likelihood (REML), with degrees of freedom estimated using the residual method. Effect estimate (denoted as β), 95% confidence interval (CI), and P values were reported. Analysis of variance (ANOVA) was used to compare the cortical CBF indices across varying PLDs. F-statistics and degrees of freedom (effect and error) were reported for ANOVA. Measurement values were presented as mean ± standard deviation. A P value of <0.05 was considered statistically significant.

## 3. Results

### 3.1 Hyperintensity in the CBF maps

In previous studies [5, 16, 20], a post-labeling delay (PLD) of 300 ms was used for CBF mapping to match the short bolus arrival time observed in mice under isoflurane anesthesia. Accordingly, we selected the pCASL images acquired at a PLD of 300 ms as the representative dataset for examining hyperintense voxels.

A total of 12 axial slices covering the mouse brain were acquired, as demonstrated by the control image of pCASL MRI (Figure 1A). In the averaged CBF index maps (N = 10), Voxels with high signal intensity showing clear contrast from surrounding brain tissue were observed across the brain, particularly in the occipital lobe (Slice #2–4; Figure 1B). A histogram was computed across brain voxels (Figure 1C) and approximated a Gaussian distribution. Fitting a Gaussian function 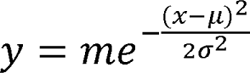, where m is the amplitude, µ the center, and a the standard deviation, yielded µ = 3.412% and a = 0.737%. Comparing the histogram with the fitted curve revealed a positive asymmetry, with an upper-tail excess corresponding to hyperintense voxels in the pCASL image. Considering regional variations in CBF, we defined hyperintense voxels in the pCASL image as those exceeding the threshold of μ + 3σ (i.e., 5.623%) in terms of ΔM/M_0_ signals.

**Figure 1.**
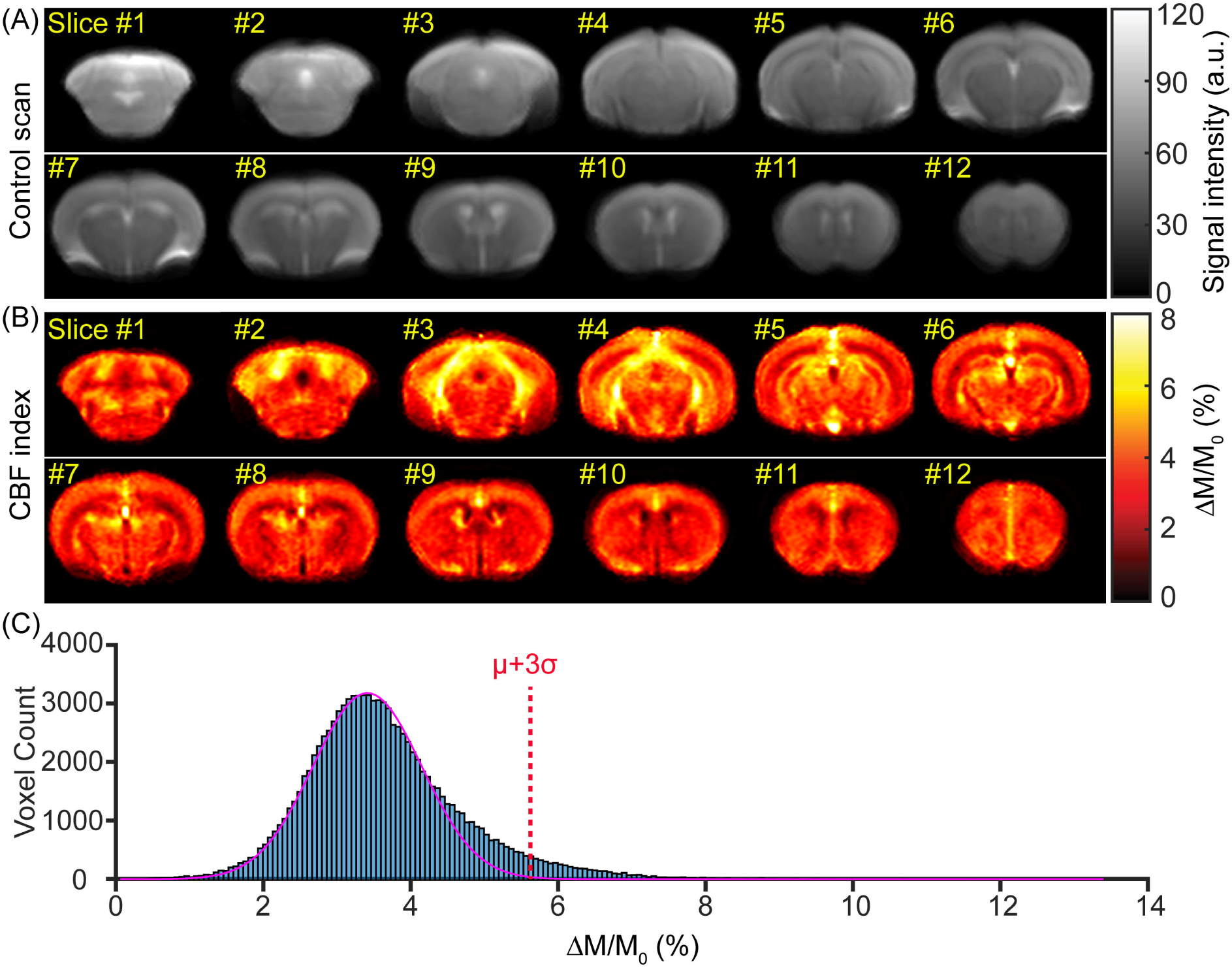
Histogram analysis of pseudo-continuous arterial spin labeling (pCASL) signals (N = 10). (A) Mean pCASL control images. (B) Mean cerebral blood flow (CBF) index maps. (C) Histogram of pCASL CBF indices; the magenta line shows the fitted Gaussian function, and the red dotted line marks the threshold at µ+ 3a.

Using the determined threshold, we extracted hyperintense matrices representing the spatial distribution of hyperintense voxels across the mouse brain (Figure 2A). By comparing TOF angiography with the vasculature atlas [19], we identified major arteries and veins, including DCA and TS in Slice #2; PCA and CS in Slice #3; VG and PCA in Slice #4; VG and PComA in Slice #5; TSV in Slice #6-7; azPA in Slice #8-10; ACA in Slice #10-11; and SSS in Slice #11-12 (Figure 2B). RARE images at TE = 150 ms showed excellent suppression of tissue signal (theoretically ∼1.931% residual given tissue T_2_ ≈ 38 ms at 11.7T [5]) and highlighted CSF in V4, AQ, V3, and LV (Figure 2C). Combining hyperintense, vascular, and CSF signals into a pseudo-color composite (red/green/blue, respectively) revealed broad co-localization—hyperintense voxels with vessels (shown in yellow) and with ventricles (shown in purple), indicated by arrows in Figure 2D. In human pCASL, hyperintense voxels arise predominantly in arteries; accordingly, pCASL hyperintensity is often termed the arterial-transit artifact [1]. In mice, however, hyperintense voxels extend beyond arteries to major draining veins and even ventricular structures (e.g., V3 and LV), reflecting a distinct distribution pattern.

**Figure 2.**
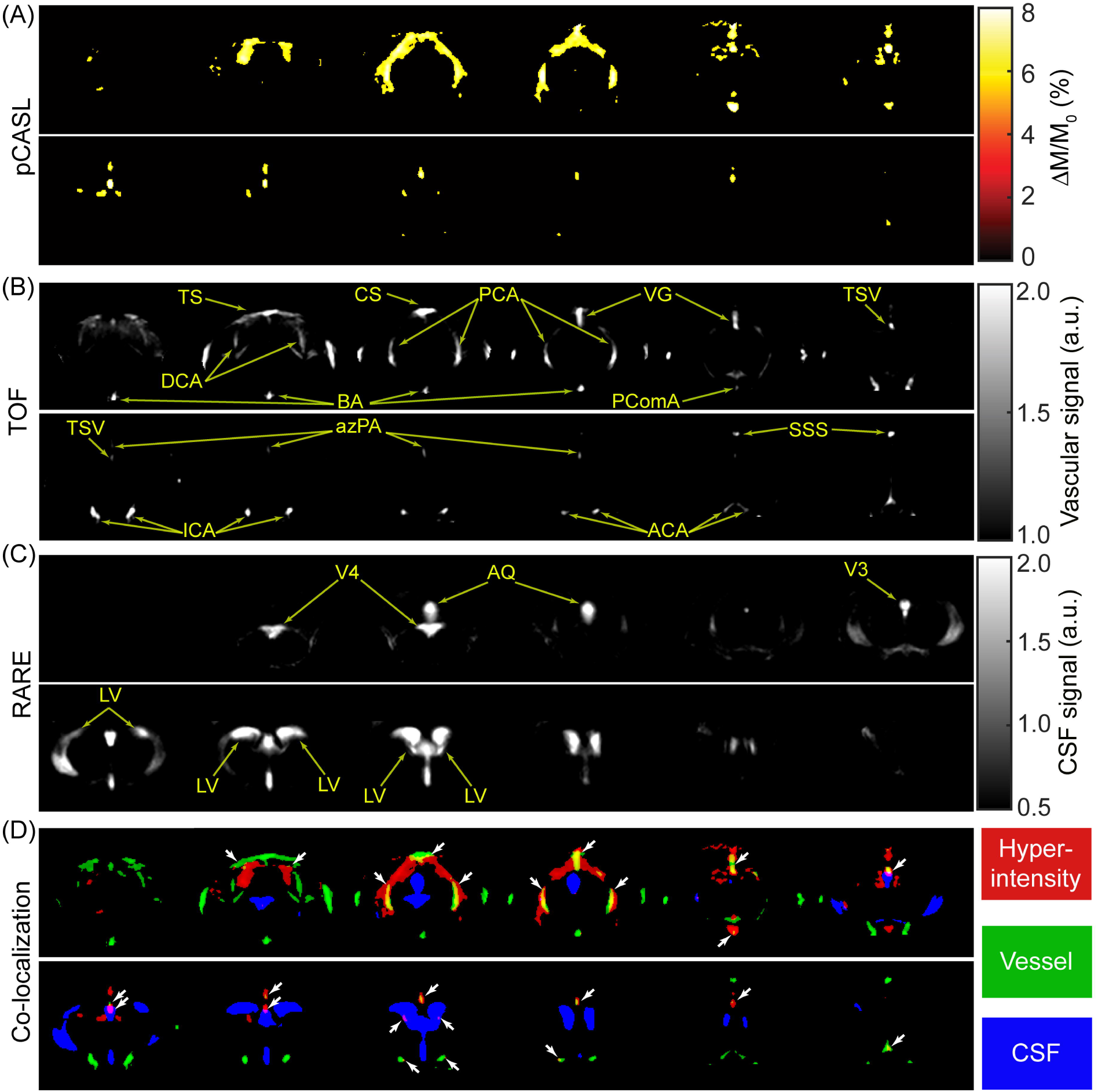
Co-localization of hyperintense voxels with vessels and ventricles (N = 20). (A) CBF index maps showing the spatial distribution of hyperintense voxels (n = 10). (B) Mean time-of-flight (TOF) angiography (n = 10). Abbreviations: DCA, dorsal cerebellar artery; TS, transverse sinus; CS, confluence of sinuses; BA, basilar artery; PCA, posterior cerebral artery; VG, vein of Galen; PComA, posterior communicating artery; TSV, thalamostriate vein; ICA, internal carotid artery; azPA, azygos pericallosal artery; ACA, anterior cerebral artery; SSS, superior sagittal sinus. (C) Mean cerebrospinal fluid (CSF) images acquired with RARE MRI at TE = 150 ms (n = 10). Abbreviations: V4, fourth ventricle; AQ, cerebral aqueduct; V3, third ventricle; LV, lateral ventricle. (D) Pseudo-color composite illustrating co-localization of hyperintense voxels with vessels (yellow) and ventricles (purple); arrows indicate representative co-localized sites.

### 3.2 pCASL scans in varying PLD values

Clear hyperintense voxels were observed across PLD values (Figure 3), particularly at shorter PLDs of 25–500 ms (Figures 3A-3J). At longer PLDs of 1000–2000 ms (Figures 3M-3P), these hyperintense voxels appeared diminished, primarily due to signal decay and the use of a suboptimal display range. When the display range was halved (0–4%), hyperintense voxels evident at shorter PLDs still showed stronger intensities than the surrounding tissue (Supporting Figure S1).

**Figure 3.**
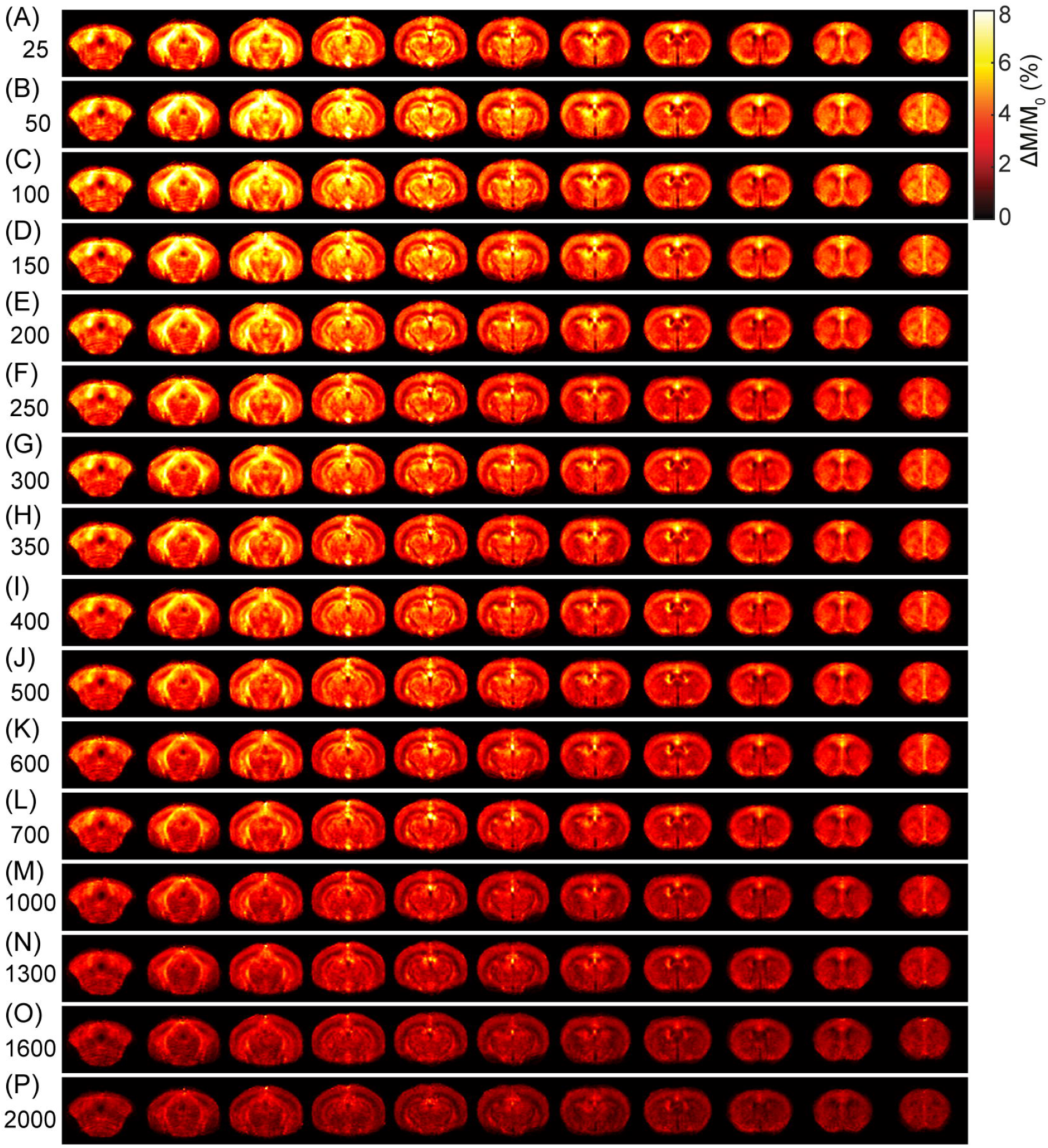
Mean CBF index maps across post-labeling delays (PLDs) (N = 10). Panels show PLD = 25 – 2000 ms: (A) 25, (B) 50, (C) 100, (D) 150, (E) 200, (F) 250, (G) 300, (H) 350, (I) 400, (J) 500, (K) 600, (L) 700, (M) 1000, (N) 1300, (O) 1600, (P) 2000. A fixed color scale is used across panels.

Regional analyses were conducted across four categories of ROIs: tissue (cortex), arteries (DCA, PCA, PComA, azPA, ACA), veins (TS, CS, VG, TSV, SSS), and ventricles (LV) (Figure 4A). The CBF indices (ΔM/M_0_) of all regions exhibited a significant dependence on PLD (β ≤ - 1.477 %/ms, P < 0.001, Figures 3B-3M), consistent with the T_1_-related decay of labeled spins during transit and exchange. Relative to cortex, the linear mixed-effects model showed significant region-by-PLD interaction effects for DCA (β = −1.236 %/s, 95% CI [−1.875, −0.597], P < 0.001), PCA (β = −1.473 %/s, 95% CI [−2.111, −0.834], P < 0.001), azPA (β = −1.266 %/s, 95% CI [−1.905, −0.627], P < 0.001), PComA (β = −2.312 %/s, 95% CI [−2.951, −1.673], P < 0.001), TS (β = −0.913 %/s, 95% CI [−1.552, −0.275], P = 0.005), CS (β = −0.758 %/s, 95% CI [−1.397, −0.119], P = 0.020), VG (β = −0.970 %/s, 95% CI [−1.609, −0.332], P = 0.003), TSV (β = −1.674 %/s, 95% CI [−2.312, −1.035], P < 0.001), and SSS (β = −0.759 %/s, 95% CI [−1.397, −0.120], P = 0.019), but not for ACA (β = −0.458 %/s, 95% CI [−1.097, 0.181], P = 0.159) or LV (β = −0.400 %/s, 95% CI [−1.039, 0.238], P = 0.219). These negative interaction terms indicate that major hyperintense regions exhibited faster signal decay with increasing PLD than cortical tissue, consistent with their intravascular origin.

**Figure 4.**
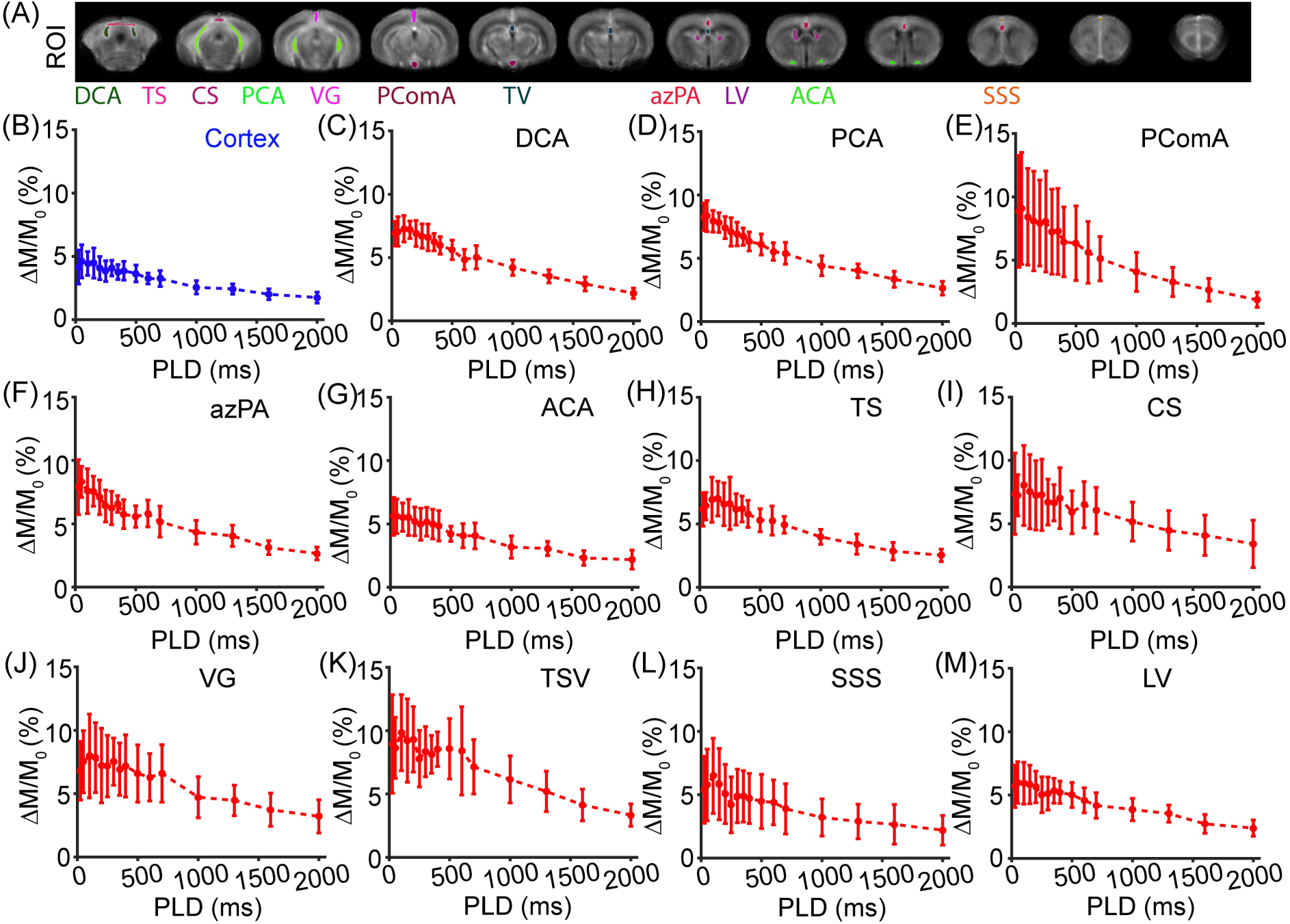
CBF indices across cerebral regions. (A) Regions of interest (ROIs) overlaid on CBF index maps. Four categories of ROIs were defined: tissue (cortex), arteries (DCA, PCA, PComA, azPA, ACA), veins (TS, CS, VG, TSV, SSS), and ventricles (LV). (B–M) Regional CBF indices for cortex, DCA, PCA, PComA, azPA, ACA, TS, CS, VG, TSV, SSS, and LV, respectively.

Given the detection-sensitivity constraints of pCASL MRI, we reanalyzed cortical CBF indices to identify a PLD that would not substantially compromise sensitivity. ANOVA revealed a significant PLD effect (*F*[15,144] = 12.210, P < 0.001). Using Student’s *t*-tests with Holm–Bonferroni correction, the cortical CBF index at 300 ms (an optimal PLD used in literature [16, 20]) differed significantly from values at PLD ≥ 600 ms. Accordingly, we selected a PLD of 500 ms as a pragmatic compromise between sensitivity (favoring shorter PLDs) and suppression of hyperintense voxels (favoring longer PLDs).

### 3.3 pCASL scans with varying vascular suppression gradients

Seven vascular-crushing gradient strengths were tested (Figures 4A–4G). Linear mixed-effects modeling showed a significant dependence of CBF index on gradient strength for cortex (β = −0.022 %·cm/Gauss, 95% CI [−0.034, −0.010], P < 0.001; Figure 5H), DCA (β = −0.046, 95% CI [−0.063, −0.029], P < 0.001; Figure 5I), PCA (β = −0.050, 95% CI [−0.073, −0.027], P < 0.001; Figure 5J), azPA (β = −0.050, 95% CI [−0.084, −0.016], P = 0.005; Figure 5L), ACA (β = −0.038, 95% CI [−0.058, −0.018], P < 0.001; Figure 5M), TS (β = −0.061, 95% CI [−0.094, −0.027], P < 0.001; Figure 5N), CS (β = −0.050, 95% CI [−0.091, −0.008], P = 0.021; Figure 5O), VG (β = −0.113, 95% CI [−0.175, −0.051], P < 0.001; Figure 5P), and TSV (β = −0.071, 95% CI [−0.113, −0.030], P = 0.002; Figure 5Q). There were decreasing trends in CBF index for PComA (β = - 0.018, 95% CI [−0.038, 0.002], P = 0.076; Figure 5K), SSS (β = −0.088, 95% CI [−0.186, 0.011], P = 0.078; Figure 5R), and LV (β = −0.022, 95% CI [−0.044, 0.001], P = 0.062; Figure 5S), but did not reach statistical significance. The negative effect estimates indicate progressive signal reduction with stronger gradients over the observed regions, consistent with diffusion- and flow-related dephasing.

**Figure 5.**
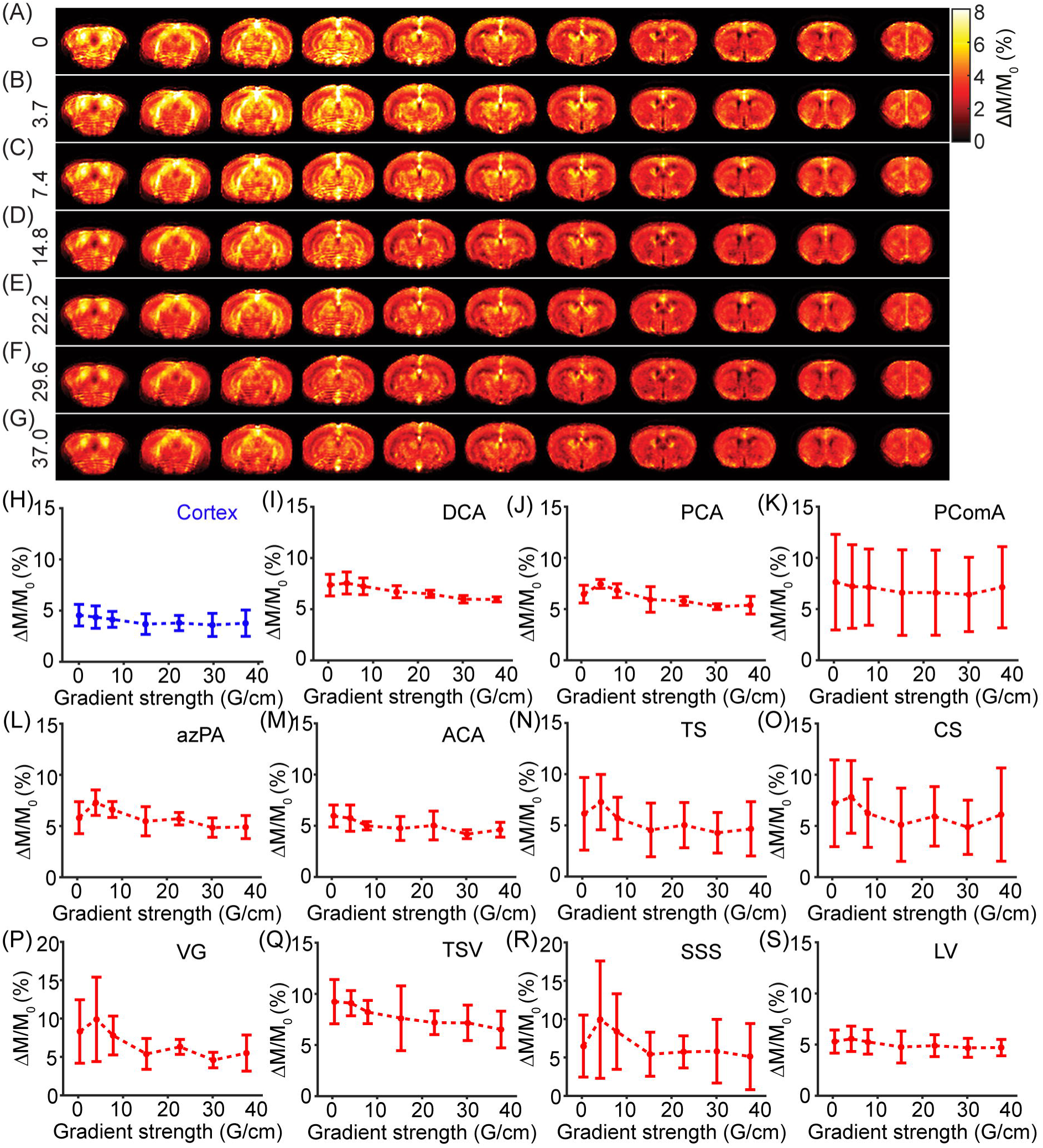
Mean CBF index maps and regional CBF indices across varying vascular-crushing gradients (N = 4). (A – G) Mean CBF index maps with triple-axis crusher gradient strengths of 0, 3.7, 7.4, 14.8, 22.2, 29.6, and 37.0 G/cm, respectively. Corresponding velocity encoding (VENC) values were ∞, 12.4, 6.2, 3.1, 2.1, 1.6, and 1.2 cm/s. (H – S) Regional CBF indices for cortex, DCA, PCA, PComA, azPA, ACA, TS, CS, VG, TSV, SSS, and LV, respectively.

Relative to cortex, a significant region-by-gradient-strength interaction was detected only for VG (β = −0.091, 95% CI [−0.177, −0.041], P = 0.041), with no significant interactions in other ROIs (P ≥ 0.140), indicating that increasing vascular-crushing strength did not preferentially suppress most hyperintense regions compared with cortex. For example, even at 37 G/cm, hyperintensities in DCA, PCA, TS, and CS remained prominent (Figure 5G).

To investigate the cause of unsatisfactory vascular suppression, we performed numerical simulations assuming laminar arterial flow (Figure 6A). Under vascular-crushing gradients, spins accrue velocity-dependent phase, producing heterogeneous phase across the arterial cross-section and consequent signal modulation (Figure 6B). When spins with different phases are sampled within the same voxel, partial cancellation occurs. Residual peak intensity within the flow cross-section decreased monotonically with increasing acquisition voxel size (Figure 6C), and this pattern held across a range of arterial diameters (Figure 6D). Defining the “optimal voxel size” for each diameter as the minimal voxel size yielding 95% vascular suppression (i.e., 5% residual peak intensity), we observed a strong linear relationship between artery diameter and optimal voxel size (*y* = 0.777*x* − 13.897, R² = 0.987, P < 0.001; Figure 6E), indicating that voxel sizes on the order of, or larger than, the arterial diameter are required for robust suppression. This implies a trade-off between spatial resolution and effective vascular suppression with crusher gradients. We further examined flow-profile effects and found that the laminar fraction significantly reduced the residual peak intensity, with a significant interaction with artery diameter (β = −0.003 %/µm, 95% CI [−0.003, −0.003], P < 0.001; Figure 6F); higher laminar fractions yielded stronger suppression than plug flow. Experimentally, our maximum crusher strength was 37.0 G/cm (50% of the system limit on the current 11.7 T scanner). Simulations extending gradient strength beyond 50% demonstrated a significant gradient-strength-by-voxel-size interaction (β = 0.223 %/Gauss, 95% CI [0.166, 0.279], P < 0.001; Figure 6G). As shown in Figure 6G, increasing gradient strength provides little additional suppression at small voxel sizes (left of the heat map), whereas at larger voxel sizes strong crushers are unnecessary (right of the heat map). The hyperintense ROIs in this study included major cerebral arteries and veins in mice, which have relatively large calibers (e.g., posterior cerebral artery: 509 ± 83 μm). By contrast, the pCASL acquisition used an in-plane voxel size of 156 × 156 μm², far smaller than the optimal voxel size predicted for effective vascular suppression.

**Figure 6.** Simulations of the effects of artery diameter, voxel size, and vascular-crushing gradient strength. (A) Velocity map of laminar flow with a peak velocity of 10 cm/s, artery diameter of 300 µm, and simulated field of view of 1000 µm. (B) Real and imaginary components of flow signals after incorporating flow-dependent phase shifts under vascular-crushing gradients. (C) Simulated laminar-flow signals for varying voxel sizes (1 × 1 µm² to 300 × 300 µm²). (D) Dependence of peak signal intensity on voxel size and artery diameter. (E) Relationship between artery diameter and the optimal voxel size required to achieve 95% vascular signal suppression. Here, “optimal voxel size” refers to the minimal voxel dimension that yields 95% vascular signal suppression. (F) Dependence of peak signal intensity on artery diameter (100 – 400 µm) and laminar fraction (0 – 1). Laminar fraction denotes the proportion of laminar flow relative to plug flow; for example, 0.8 indicates 80% laminar and 20% plug flows. (G) Dependence of peak signal intensity on voxel size (1–300 µm) and vascular-crushing gradient strength (0 – 70 G/cm).

## 4. Discussion and conclusions

We characterized hyperintensity in mouse pCASL-based CBF maps as a function of PLD and vascular-crushing gradient strength. A PLD of 500 ms provided a tradeoff between detection sensitivity and suppression of hyperintense voxels. By contrast, increasing crusher-gradient strength yielded little additional suppression for the hyperintense signals.

CBF is a potential biomarker for monitoring vascular pathology, assessing therapeutic efficacy, and evaluating physiological perturbations following drug administration [7, 21, 22]. Evaluating CBF in diseases characterized by vascular dysfunction can aid both diagnosis and prognosis [23, 24]. For example, perfusion imaging helps delineate hypoperfusion boundaries in ischemic stroke, facilitating the identification of the core region and ischemic penumbra [25]. Additionally, early-stage CBF recovery after cardiac arrest serves as a predictor of neurological outcomes [7]. Due to neurovascular coupling, CBF alterations occur not only in vascular pathologies but also in metabolic disorders such as Alzheimer’s disease [26, 27]. Furthermore, CBF can be integrated into multiparametric studies alongside other imaging and physiological techniques to comprehensively characterize pathophysiological changes [28–30]. In particular, combining vascular physiology with pathological assessments [31, 32] enhances our understanding of the full spectrum of pathophysiological changes [33–35].

The water extraction fraction—the proportion of arterial water extracted by tissue—is markedly lower in mice (59.9 ± 3.2%) [17] than in humans (95.5 ± 1.1%) [36]. As a result, a substantial fraction of labeled water bypasses capillary exchange and drains directly into the venous system, producing venous hyperintensities. Accordingly, hyperintense voxels in human pCASL are predominantly arterial (arterial-transit artifact) [1], whereas in mice they extend to major draining veins—the superior sagittal sinus, transverse sinus, confluence of sinuses, great vein of Galen, and thalamostriate vein. This arterial-and-venous distribution complicates PLD selection: a PLD long enough to attenuate arterial signal may still leave venous hyperintensity, while excessively long PLDs erode sensitivity. We therefore adopted a pragmatic criterion that prioritizes tissue perfusion signal while accommodating long transit times; under isoflurane, a PLD of 500 ms provided a reasonable balance for mouse pCASL.

Apart from the arterial and venous hyperintensities, hyperintense voxels were observed in the lateral ventricles, suggesting that the used experimental parameters (labeling duration of 1800 ms) allow the visualization of pCASL signals delivered to ventricles through choroid plexuses. However, the current ROI-based ventricular data were not suitable for estimating ventricular bolus arrival time owing to parenchymal partial-volume contamination, as indicated by the absence of the biphasic (rise-then-fall) slope predicted by the Buxton’s kinetic model [2]. Future work could use a long-TE readout (to further attenuate tissue signal) and more selective ventricular masks to enable estimation of ventricular arrival time when relevant.

Bipolar crusher gradients are widely used to suppress intravascular signal [1, 10, 11, 16]. In our data, however, increasing crusher strength did not consistently reduce hyperintensity, primarily because the small voxel size relative to arterial caliber limits intravoxel phase dispersion. Although high spatial resolution (i.e., small voxels) is desirable for resolving fine structures, it reduces intravoxel dephasing and thus the potential for intravoxel signal cancellation. Consequently, pCASL hyperintensities in major arteries and veins in mice are unlikely to be eliminated by stronger crushers. Given the associated signal-to-noise penalty (via TE prolongation and diffusion weighting), the additional use of crusher gradients for vascular suppression is not recommended for mouse pCASL MRI. Differences in the efficacy of vascular crushing between humans and mice reflect interspecies vascular scaling. Although murine vessels are absolutely smaller, their calibers occupy a larger fraction of the mouse brain—and of typical preclinical voxels—than in humans, reducing intravoxel phase dispersion and limiting crusher effectiveness.

Diverse hyperintensity behaviors in pCASL-based CBF maps (human vs. mouse) across varying post-labeling delays and vascular-crushing gradients necessitate dedicated optimization studies for mouse pCASL. Direct back-translation of human pCASL knowledge is unlikely to account fully for interspecies differences and the experimental differences between preclinical and clinical settings. Unlike in human pCASL, crusher gradients are not a practical optimization strategy in mice, and future mouse pCASL studies should instead prioritize PLD selection.

Findings in this study should be interpreted within the context of the following limitations. First, all experiments were conducted under isoflurane, which is vasoactive and increases baseline CBF in a dose-dependent manner [37]. Therefore, the optimal PLD identified here may not generalize to conditions without anesthesia or other anesthesia agents (for example, dexmedetomidine). Second, analyses on lateral ventricles were based on relatively large ROIs and were affected by parenchymal partial-volume contamination, precluding kinetic fitting for estimating ventricular bolus arrival time. Third, the sample size for the crusher-gradient optimization study was modest and may have reduced sensitivity to small regional effects. While this is a limitation, the alignment between experimental findings and simulations supports the robustness of the overall conclusions. The present study provides a helpful guidance for future methodological refinement. Follow-up work evaluating alternative anesthetic protocols and larger cohorts will further validate and generalize these findings.

In summary, under isoflurane anesthesia in mice, hyperintense voxels in pCASL-based CBF maps are not confined to arteries but extend to major draining veins. A post-labeling delay of 500 ms offers a practical trade-off between detection sensitivity and suppression of hyperintensity voxels. Increasing vascular-crushing gradient strength provided marginal additional suppression and is therefore not recommended for routine use.

## Author Contributions Statement

Xiuli Yang: Methodology; Investigation; Validation; Writing – Original Draft Preparation. Yuguo Li & Adnan Bibic: Investigation; Writing – Review & Editing. Zhiliang Wei: Conceptualization; Methodology; Software; Resources; Data Curation; Funding Acquisition; Writing – Original Draft Preparation; Writing – Review & Editing.

## Data Availability Statement

Data will be available upon request to the authors.

## Acknowledgements

This work was supported by the Grant Sponsors: National Institutes of Health (NIH) R01 AG081932 and NIH P41 EB031771.

## Supporting Information

**Figure S1** CBF maps of long PLDs: (A) 1000 ms, (B) 1300 ms, (C) 1600 ms, and (D) 2000 ms (N = 10). CBF index defined by the ratio between difference signal and equilibrium magnetization was displayed at the range of 0-4%.

