## Supplemental figure for "Hyperintense signals in cerebral blood flow maps acquired with pseudo-continuous arterial spin labeling MRI in mice"

**Supporting Information**

1. CBF index map of long post-labeling delays (PLDs)

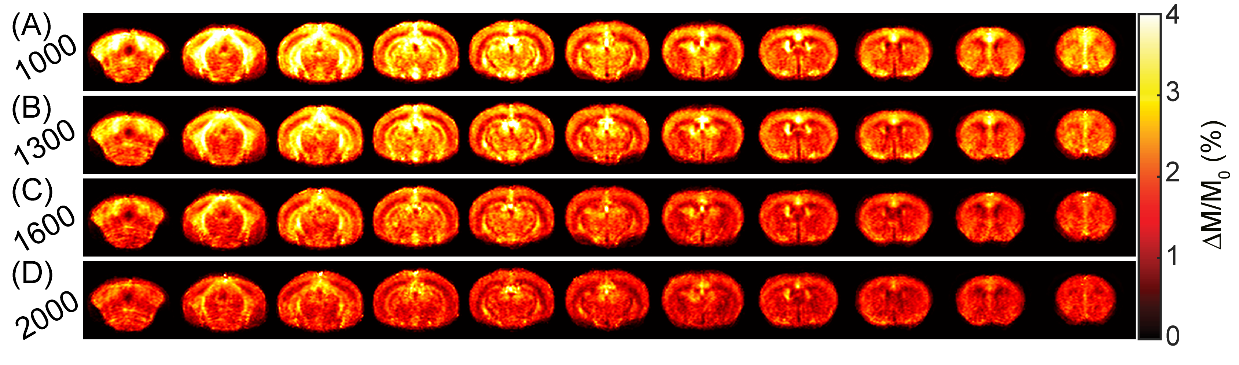

**Figure S1** CBF maps of long PLDs: (A) 1000 ms, (B) 1300 ms, (C) 1600 ms, and (D) 2000 ms (N = 10). CBF index defined by the ratio between difference signal and equilibrium magnetization was displayed at the range of 0-4%.
